# Epileptic Spike Detection Using Neural Networks with Linear-Phase Convolutions

**DOI:** 10.1101/2020.10.08.330936

**Authors:** Kosuke Fukumori, Noboru Yoshida, Hidenori Sugano, Madoka Nakajima, Toshihisa Tanaka

## Abstract

To cope with the lack of highly skilled professionals, machine learning with proper signal processing is key for establishing automated diagnostic-aid technologies with which to conduct epileptic electroencephalogram (EEG) testing. In particular, frequency filtering with the appropriate passbands is essential for enhancing the biomarkers—such as epileptic spike waves—that are noted in the EEG. This paper introduces a novel class of neural networks (NNs) that have a bank of linear-phase finite impulse response filters at the first layer as a preprocessor that can behave as bandpass filters that extract biomarkers without destroying waveforms because of a linear-phase condition. Besides, the parameters of the filters are also data-driven. The proposed NNs were trained with a large amount of clinical EEG data, including 15,833 epileptic spike waveforms recorded from 50 patients, and their labels were annotated by specialists. In the experiments, we compared three scenarios for the first layer: no preprocessing, discrete wavelet transform, and the proposed data-driven filters. The experimental results show that the trained data-driven filter bank with supervised learning behaves like multiple bandpass filters. In particular, the trained filter passed a frequency band of approximately 10–30 Hz. Moreover, the proposed method detected epileptic spikes, with the area under the receiver operating characteristic curve of 0.967 in the mean of 50 intersubject validations.

## I. Introduction

EPILEPSY is a neurological disorder that is said to affect 50 million patients worldwide. In particular, childhood epilepsy affects an individual’s cognitive activity. Early appropriate diagnosis helps patients reduce future brain damage. In the diagnosis, the measurement of an electroencephalogram (EEG) along with a medical examination is essential for determining the type of seizure symptom. The examination requires clinical knowledge and experience, but epilepsy specialists with these skills are in chronically short supply. This has motivated the development of an automated diagnostic tool to support epileptologists.

One of the essential biomarkers in diagnosing epilepsy is an epileptic spike called a paroxysmal discharge, which is frequently present in a patient’s interictal EEG [1]. To support the detection of epileptic spikes, several automated detection approaches are making great advances. To implement the automatic detection of epileptic spikes, supervised learning is one effective method. To efficiently train the machine learning models, the EEG signal is generally decomposed into standard clinical frequency bands of interest—such as *δ*, *θ*, *α*, *β*, and *γ*—before the learning [2]. While conducting such training, it is necessary to select the frequency bands appropriately, which depends on several factors, such as the EEG measurement method, measurement environment, the type of epilepsy, and epileptologists’ skills. However, in various studies, a range of frequencies or frequency bands of interest has been empirically selected. Douget et al. [3] used discrete wavelet transform (DWT) to obtain a set of subbands with a range of 4–32 Hz for the analysis of both intracranial and scalp EEG. Carey et al. [4] used an infinite impulse response Butterworth bandpass filter with a frequency band of 1–30 Hz. In addition, Maurice et al. [5] employed 0.5–70 Hz band-pass filter with a third-order Butterworth and 60 Hz notch filter with a fourth-order Butterworth to detect spikes from an intracranial EEG. For epileptic seizure detection, Iesmantas et al. [6] used seven bandpass filters to divide the EEG into seven frequency bands of <4 Hz, 4–7 Hz, 7–13 Hz, 13–15 Hz, 14–30 Hz, 30–45 Hz, and 45–70 Hz.

With the advent of deep neural networks, model parameters could learn from observed data, including feature extraction methods. In particular, convolutional neural networks (CNN) can extract features by applying filters to input data [7], [8]. However, each filter in the convolutional layer has a high degree of freedom, even though the predefined filters in previous studies have been designed with a linear-phase constraint to preserve the waveform shape. It is undesirable to destroy the waveform shape using unconstrained filtering, because the manual identification of epileptic biomarkers by clinical experts is also an essential requirement of the diagnosis.

This paper hypothesizes that the frequency subbands can be estimated on the basis of the data from an epileptic EEG labeled by clinical specialists. To this end, we propose the use of supervised learning to find filter coefficients regarded as a one-dimensional (1D) convolutional layer under a linear-phase constraint. This layer can be connected to general neural networks, such as CNN and artificial neural networks (ANN), as a classifier. Furthermore, because no dataset of epileptic spike detection is, to the best of our knowledge, publicly available, this paper built a large medical dataset—containing 15,833 epileptic spikes, 15,004 nonepileptic discharges, and the corresponding 30,837 labels—to train the parameters in the proposed model.

## II. Related Work

### A. Feature Extraction

Recently, many studies of epileptic EEGs have applied signal decomposition methods using DWT in a preprocessing stage [3], [9]–[15]. However, in these cases, the parameter selection frequency range of the bandpass filters is empirically given.

Cheong et al. [16] used DWT to decompose the signal into frequency subbands from the delta band to the gamma band (0–63 Hz). Gutierrez et al. [9] applied a bandpass filter in the range of 0.5–70 Hz. Then, they obtained wavelet coefficients from the filtered signal to classify epileptic spikes. Similarly, a range of 0.5–70 Hz was extracted with a Butterworth filter to obtain wavelet coefficients [15].

Other studies have utilized narrow bandpass filter ranges for preprocessing. Polat et al. [17] applied a bandpass filter range of 0.53–40 Hz and then used the discrete Fourier transform to extract the features for the decision tree classifier. Khan et al. [12] used the range of 0–32 Hz decomposed by DWT because most of epileptic information lies in the range of 0.5–30 Hz. Similarly, Douget et al. [3] and Indiradevi et al. [18] adopted DWT with Daubechies 4 (DB4) to extract the frequency band of 4–32 Hz. Moreover, Fergus et al. [19] used the range of only 0–25 Hz, although they did not use DWT but a Butterworth filter. Thereafter, they employed the holdout technique and k-fold cross-validation, passing into many different classifier models for distinguishing seizure and nonseizure EEG records.

In these studies for the classification or detection of epilepsy, DWT decomposition and other filtering methods were effective. As seen above, although the frequency range, including the epileptic information, is roughly known to be less than about 60 Hz, the selection of cut-off frequencies depends on several factors, such as the designer of the automated system, the type of epilepsy, the epileptologists on diagnosis, and so forth. This motivated us to identify the filter parameters based on data.

### B. Convolutional Neural Networks

A type of neural network (NN) that demonstrates excellent performance—especially in the field of image or video recognition [20], [21]—is the CNN. The CNN is an extended NN that has an input layer, multiple hidden layers, and output layer. In general, the hidden layers consist of convolutional layers, and a fully connected layer is used as the output layer. The convolution layer applies a convolution to the input and forwards the result to the next layer. Let *X* = {*x*_0_, *x*_1_, … , *x*_*N*−1_}, *Y* = {*y*_0_, *y*_1_, … , *y*_*M*−1_}, and *H* = *h*_0_, *h*_1_, … , *h*_*L*−1_ be a 1D input signal, a 1D output signal, and a convolutional kernel, where *N*, *M*, and *L* are the length of *X*, *Y*, and *H*, respectively. For the sake of simplicity, *L* is assumed to be even. Focusing on one layer, the input *X* is convolved with the kernel *H*, and the output *Y* is generated as follows:

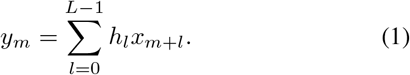

The flattened layer smoothes multiple convolved signals into a single dimension. Then, the fully connected layer multiplies all input neurons by their weight coefficients and connects them to the output.

Some recent studies have applied a CNN-based model to EEG signals [7], [8], [22], [23]. Ullah et al. [7] used 1D convolution to extract features by filtering time series EEG. Zhou et al. [23] directly input both of the multichannel time series EEG signals and their frequency domain signals into a CNN-based model. Such studies using CNN to detect epileptic seizures or epileptic spikes have been gaining interest.

### C. Dataset of Other Works

This section summarizes datasets of recent studies of epileptic spike detection. The most common task is the classification of epileptic spike waveforms and nonepileptic waveforms. Table I summarizes the datasets from similar studies. It should be emphasized that the dataset constructed in this paper achieved a much larger dataset (15,833 epileptic spike waveforms from 50 patients) than previous studies, in which the largest dataset in terms of spike waveforms consisted of 7,500 samples [22] and the largest one in terms of patients consisted of 50 patients [24]. Note that neither of the datasets from the previous studies is publicly available.

**TABLE I.** summary of the datasets on epileptic spike detection in other studies

| Reference and publication year | #Labels annotated as epileptic spikes | #Patients |
| --- | --- | --- |
| Wilson et al. [24], 1999 | 2,400 | 50 |
| Indiradevi et al. [18], 2008 | 684 | 22 |
| Liu et al. [25], 2013 | 142 | 12 |
| Johansen et al. [22], 2016 | 7,500 | 5 |
| Douget et al. [3], 2017 | 2,157 | 17 |
| Xuyen et al. [14], 2018 | 1,491 | 19 |
| Thanh et al. [15], 2020 | 1,442 | 17 |
| This paper | 15,833 | 50 |

## III. Method

### A. Dataset

EEG recordings were collected from 50 patients (24 males and 26 females) with childhood epilepsy with centro-temporal spikes (CECTS) [26] at the Department of Pediatrics, Juntendo University Nerima Hospital. The age range of the patients at the time of the examination was 3–12 years. The data were recorded from 16 electrodes with the international 10– 20 methods using the Nihon Koden EEG-1200 system. The sampling frequency was 500 Hz. This dataset was recorded and analyzed with the approval of the Juntendo University Hospital Ethics Committee and the Tokyo University of Agriculture and Technology Ethics Committee. Details of these EEG recordings are given in Appendix A.

First, two neurosurgeons, one pediatrician, and one clinical technologists selected a focal channel associated with the origin of the epileptic spike. In particular, CECTS is a type of focal epilepsy in which spikes appear only in a certain channel. Therefore, the annotators chose the most epileptic intense channel as the annotation channel. Peaks (minima and maxima) of the EEG at the channel were detected by a peak search function implemented with Scipy [27]. This function extracts both upward and downward peaks with a minimum distance of 100 points. Using a threshold determined at the 80th percentile value in the absolute amplitude of all peaks, meaningless peaks caused by noise, and so forth were removed. Second, the annotators labeled each peak as either an epileptic spike (spike or spike-and-wave) or nonepileptic discharge. These non-epileptic waveforms were carefully selected by the annotator from noise peaks excluding extreme voltage fluctuations caused by body movements and sweating and other possible interferences. Then, a 1-s segment was extracted at every detected peak, including 300 ms before and 700 ms after the peak. Fig. 1 illustrates an example of typical waveforms. Z-score normalization was applied with the mean value and standard deviation for each segment. It should be noted that each segment represents one candidate spike.

**Fig. 1.**
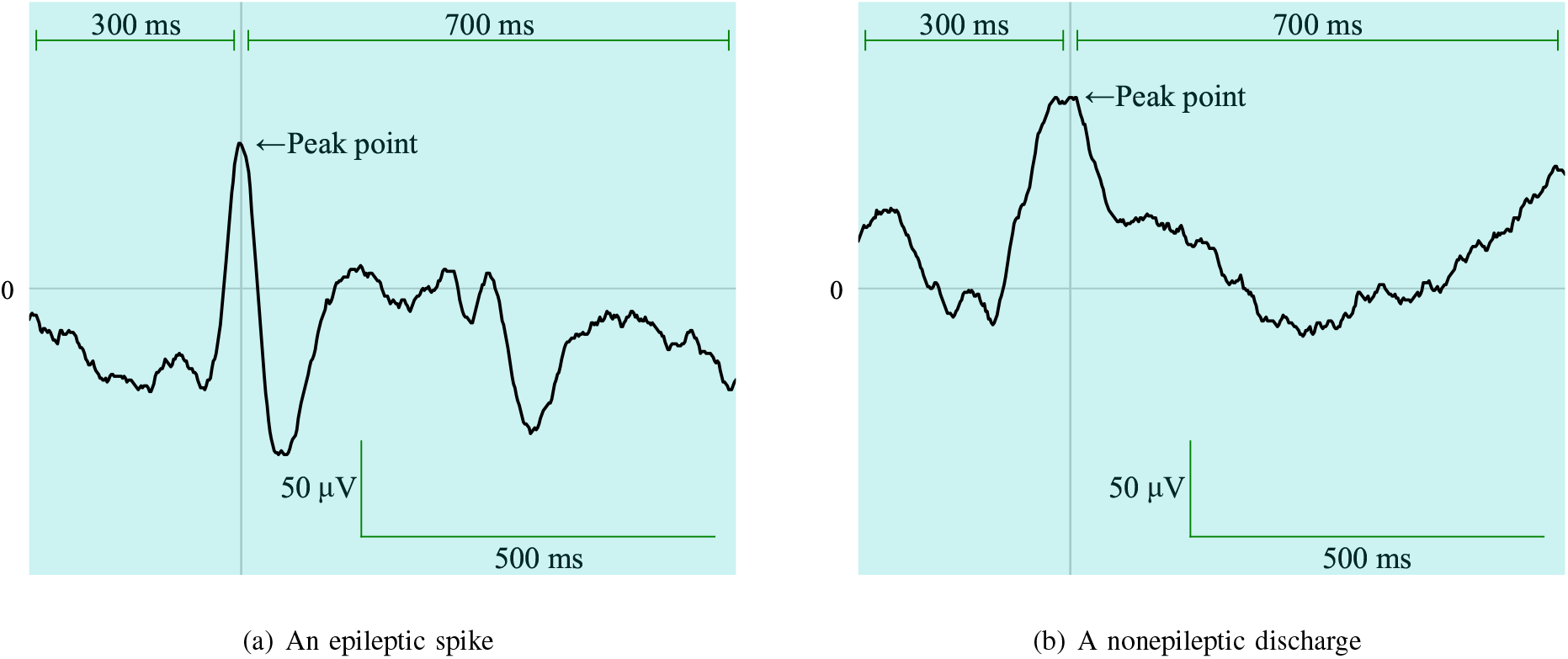
Typical waveforms of detected peaks. Each waveform is clipped into a 1-s segment.

### B. Preprocessing and Subband Decomposition

We considered two models, as shown in Fig. 2. The first model uses a predefined bank of filters. It is based on the method adopted in several previous studies. The second model involves a special convolution layer called the *linear-phase convolutional layer* (LPCL) in which the parameters are searched based on the dataset.

**Fig. 2.**
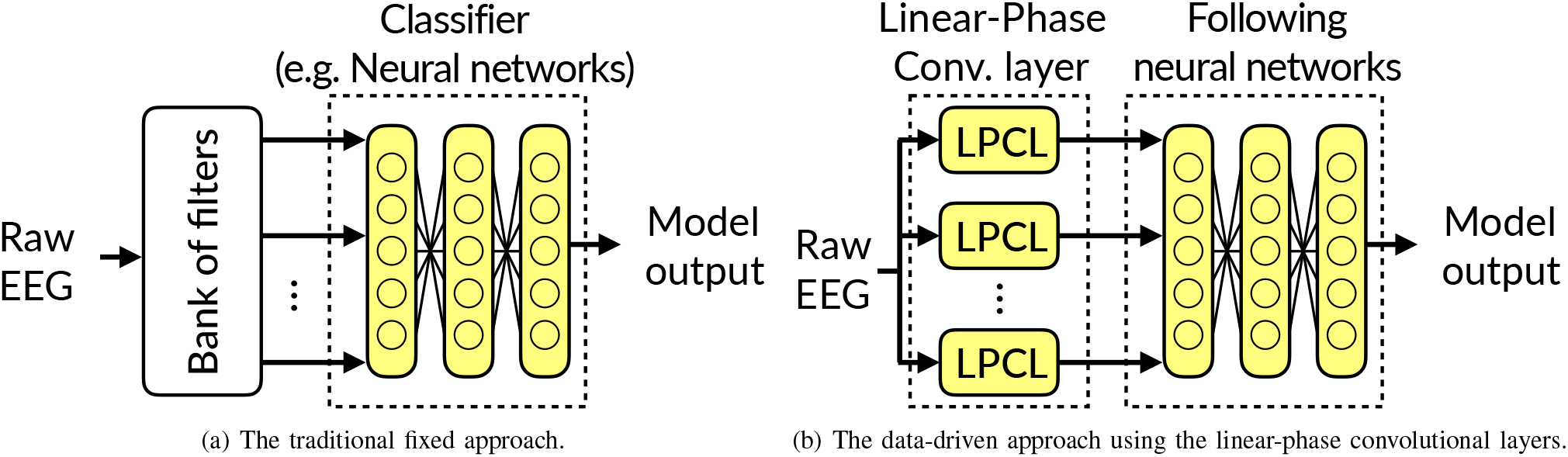
Diagrams of the two prediction models. The colored blocks contain parameters to be trained.

#### 1) Fixed approach

The first approach employs a hand-engineered preprocessing technique for each segment. DWT is applied to extract the subbands from the EEG. In this paper, the Daubechies wavelet of order 4 (DB4), which has been reported to be appropriate for analyzing EEG signals [3], [28], [29], is adopted as the mother wavelet. The input EEG is decomposed into six coefficient levels—D6, D5, D4, D3, D2, and D1—and one approximation level, A6. Then, four subbands corresponding to D6, D5, D4, or D3 are generated. Each subband represents the *θ* band (4–8 Hz), the *α* band (8–16 Hz), the *β* band (16–32 Hz), and the *γ* band (32–64 Hz), respectively [16]. The approximation level, A6, and the coefficient levels, D2 and D1, are eliminated because the low-frequency band may include breathing and eye movements. The high-frequency band can be considered noise.

#### 2) Novel data-driven approach using linear-phase convolutional layer

The convolutional layer described in Section II-B can behave as a finite impulse response (FIR) filter. However, each weight in a convolutional layer is fitted with a high degree of freedom, although FIR filters are designed with a linear-phase (LP) constraint to preserve the waveform shape. This paper proposes a convolutional layer with LP constraints, that is, the LPCL, and its implementation.

The FIR filter is realized by convolution of the discrete signal *X* = {*x*_0_, *x*_1_, … , *x*_*N*−1_} and the kernel *H* = {*h*_0_, *h*_1_, … , *h*_*L*−1_}, and the output discrete signal *Y* = *y*_0_, *y*_1_, … , *y*_*M*−1_ is calculated based on the current and past *L* − 1 inputs, much like (1). Generally, the kernel described above causes phase distortion, which can be avoided by imposing an LP constraint. When the length of the filter is even, the even symmetry and odd symmetry of the kernel yields the LP FIR filter of type-II and type-IV [30], that is:

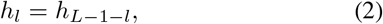

and

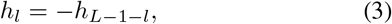

respectively. The idea behind using type-II and type-IV symmetric filters is twofold: (a) In generalizing the Haar transform to a bank of FIR filters, the multistage Haar wavelet transform is equivalent to an orthogonal matrix [31], including type-II and type-IV FIR filters with different lengths, and each filter corresponds to a bandpass filter. (b) By using type-I I and type-IV, it is possible to compose a bank of lowpass, bandpass, and highpass filters because type-II and type-IV are inherently unable to yield a highpass filter and a lowpass filter, respectively [30].

From (1) and (2), an even symmetric convolution 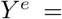 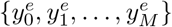 is described as follows:

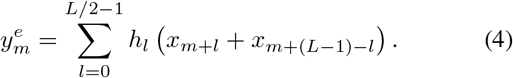

This convolution can be implemented using a lattice structure [32], as shown in Fig. 3(a). As shown in this figure, even symmetric convolution can be regarded as the product of the vector expressed by the addition of the two components in *X* and kernel *H*. This is the same operation as a weighted full connection (namely, a fully connected layer). Therefore, this can be implemented by repurposing a conventional neural network framework with the addition of *X* elements, as illustrated in Fig. 3(a). Similarly, an odd symmetric convolution 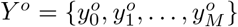 is described as follows:

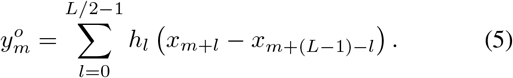

**Fig. 3.**
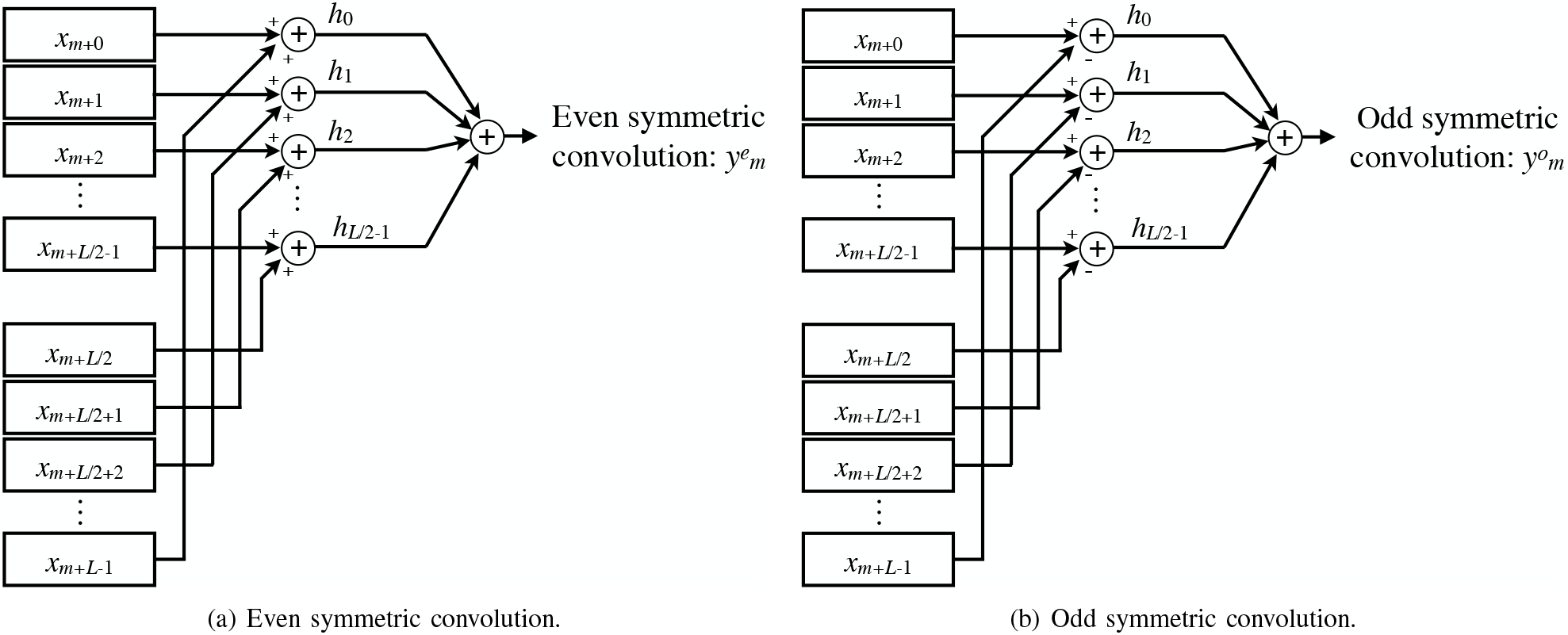
Lattice structures of the LP convolution.

Fig. 3(b) illustrates the lattice structure for (5). As this figure shows, the odd symmetric convolution can be implemented by repurposing a conventional neural network framework with the subtraction of *X* elements. These LPCLs can replace the fixed (predesigned) subband filters, as illustrated in Fig. 2. The idea is hypothesized that the LPCL can derive the frequency bands of interest from the epileptic EEG dataset.

### C. Classifier Models

Random forest (RF), ANN, and CNN are adopted as the classifiers. Although the ANN and CNN can be combined with either a traditional preprocessing technique or the proposed method, RF can be combined only with the traditional preprocessing technique.

The RF parameters are tuned using a grid search for the parameters listed in Table II. To adjust the grid search, the F1 score is used as the ranking score, and fivefold cross-validation with two subsets is used. The model architectures of the ANN and CNN are depicted in Fig. 4. To generate the initial weights of these models, the He initializer [33] is used for the layers that employ the rectified linear unit (ReLU) as the activation function. The Xavier initializer [34] is used for the other layers. These neural networks are fitted by the Adam optimizer [35] (the learning rate *η* and the scale parameters *β*_1_ and *β*_2_ are 0.001, 0.9, and 0.999, respectively) with batch size 256 while suppressing overfitting using early stopping [36].

**TABLE II.** Parameter for the random forest to be tuned by grid search

| Parameter | Candidates |
| --- | --- |
| Number of trees $N_{\text{tree}}$ | 5, 10, 20, 30, 50, 100, 300 |
| Maximum depth $D_{\text{max}}$ | 2, 4, 6, 8, 10 |

**Fig. 4.**
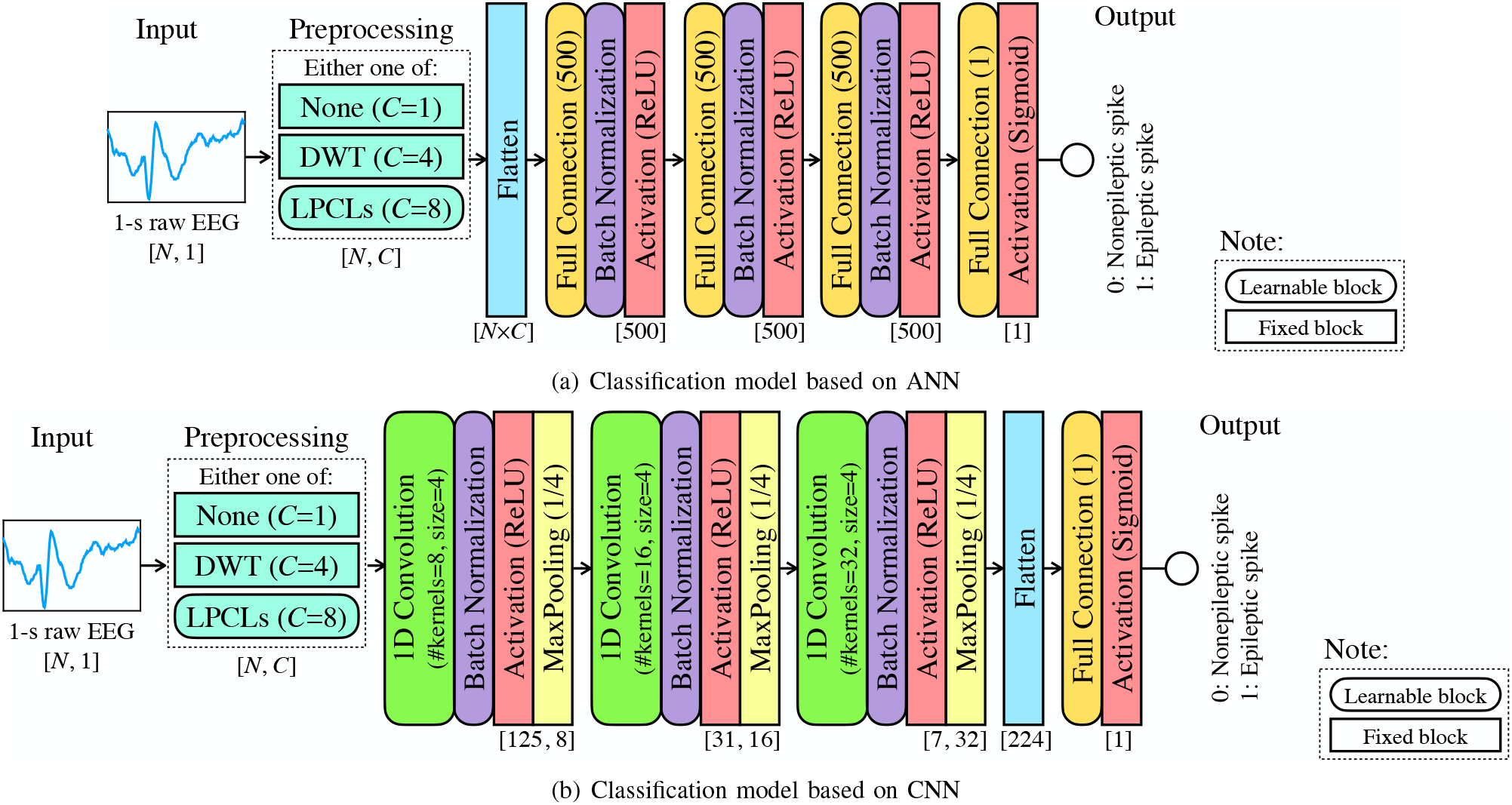
The model architectures, where *N* and *C* are the length of the input segment and the number of input subbands to the following model, respectively. When “None” is selected as the preprocessing, the raw EEG is output without any changes (*C* = 1); when “DWT,” four clinical frequency bands are extracted (*C* = 4); when “LPCLs,” the raw EEG is preprocessed by the eight LPCLs in Table III (*C* = 8). Then, the three-stacked ANN and CNN output a prediction value in the range of 0 to 1.

### D. Application of the Linear-Phase Convolutional Layer

In this paper, eight LPCLs are connected in parallel to the classification model, as illustrated in Fig. 2(b). Each LPCL setting is as shown in Table III. As in this table, there are LPCLs with different filter lengths to let the model select filters that contribute to the classification. These kernel lengths are set based on the length of the Haar transform matrix induced from the Haar wavelet [31]. That is, filter lengths of 8, 16, 32, and 64 are expected to extract the standard clinical bands of *γ*, *β*, *α*, and *θ*, respectively. At the LPCL’s filtering, the stride length is 1, and the input signal is padded with zero to keep the input and output lengths invariant. For the initialization of the coefficients in these LPCLs, the Xavier initializer [34] is used.

**TABLE III.** Settings of the LPCLs

| LPCL no. | Filter length | Constraint type |
| --- | --- | --- |
| 1 | 8 | even symmetric |
| 2 | 16 |  |
| 3 | 32 |  |
| 4 | 64 |  |
| 5 | 8 | odd symmetric |
| 6 | 16 |  |
| 7 | 32 |  |
| 8 | 64 |  |

### E. Evaluation

To validate the effectiveness of the proposed method, an experiment is performed using the dataset described in Section III-A. Recall that the classification is binary: an epileptic spike or a nonepileptic discharge. For comparison, three approaches are used: the fixed approach, the proposed data-driven approach, and an approach without preprocessing. Combining these approaches with the three classification models, a total of eight methods are compared, as shown in Table IV. In the fixed approach, a 1-s raw EEG is decomposed into four frequency bands (*θ*, *α*, *β*, and *γ* bands) using DWT. In the proposed approach, because the LPCLs act as a bank of FIR filters, a 1-s segment is input to this layer. Furthermore, in the third approach, a 1-s segment is input directly into the classification model. This approach is similar to our previous work [8].

**TABLE IV.** Methods of experimental comparison. The proposed data-driven method is combined only with the neural network models.

| Preprocessing | None | DWT | LPCL |
| --- | --- | --- | --- |
| Input feature to the following models [length, #band] | Raw EEG [500, 1] | Decomposed four frequency bands [500, 4] | Learnable eight frequency bands [500, 8] |
| Combine with RF | ✓ | ✓ |  |
| Combine with ANN | ✓ | ✓ | ✓ |
| Combine with CNN | ✓ | ✓ | ✓ |

In the experiment, intersubject validation in all combinations is performed, in which 49 patients are used as training data and the remaining patient is used for the test data. To evaluate the models, the area under the curve (AUC), F1 value, sensitivity, and specificity are employed. AUC is the area of the curve drawn by the false positive rate (FPR) and the true positive rate (TPR = Sensitivity) when the discrimination threshold is changed, and it is calculated in the following manner:

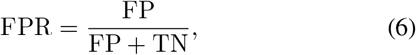

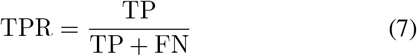

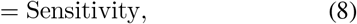

where TP, FP, FN, and TN are the numbers of a true positive, false positive, false negative, and true negative, respectively. The specificity is the true negative rate, which is calculated as follows:

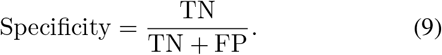

The F1 value is calculated as the harmonic mean of the precision and sensitivity. These metrics are defined as follows [37]:

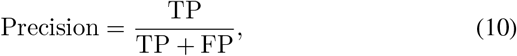

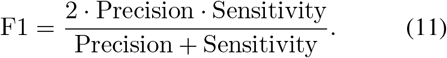

In particular, this paper employs the mean AUC and the mean F1 value (by taking 30 independent realizations) in evaluating the ANN, CNN, and LPCLs because the initial weight and initial kernel value affect the learning. In addition, because the convolution filter can be regarded as an FIR filter, the frequency response of each filter of the eight LPCLs is analyzed after training. Similar to evaluating the AUC and F1 values, the frequency response is meaned by 30 independent runs.

All experimental results are computed on a high-performance computer built with an AMD(R) EPYC(TM) 7742 CPU@2.25 GHz, 512 GB RAM, and four NVIDIA(R) A100 GPUs. The models in the experiment are constructed using Python 3.7.6 with Keras [38] and Scikit-learn [39].

## IV. Experimental Results

Table V represents the AUC, F1 value, sensitivity, and specificity by each model and preprocessing technique. This table shows the mean values of all intersubject validations. The detailed values of the intersubject validations are given in Appendix B. A statistical tests including Friedman’s oneway analysis of variance (ANOVA) [40] showed that the effects of the methods on the four metrics were significant (*F*_AUC_(1, 7) = 294, *p*_AUC_ = 7.21 × 10^−59^, *F*_F1_(1, 7) = 197, *p*_F1_ = 2.41 × 10^−38^, *F*_sen_(1, 7) = 205, *p*_sen_ = 5.89 × 10^−40^, *F*_spe_(1, 7) = 120, and *p*_spe_ = 3.10 × 10^−22^). Because the main effect of the models has been observed, a Bonferroni *post-hoc* test [40] was performed to better understand the changes in cross-correlation across the different preprocessors. Fig. 5 visualizes the numerical results and their analysis of variance of 50 intersubject validations. As shown in Fig. 5(a), significant differences in the AUC results were observed when using the preprocessors, especially for RF and ANN. Moreover, significant differences in the F1 results were observed for all classification models when using the preprocessors. In particular, the F1 results using LPCLs tended to be statistically higher than DWT in the ANN-based comparison. This is because LPCLs statistically increased specificity, as shown in Fig. 5(d). From these results, it can be seen that the preprocessing of EEG affects the classification performance, even with manually designed filters such as DWT. Furthermore, the optimal preprocessing could be learned in a data-driven method with LPCLs.

**TABLE V.**
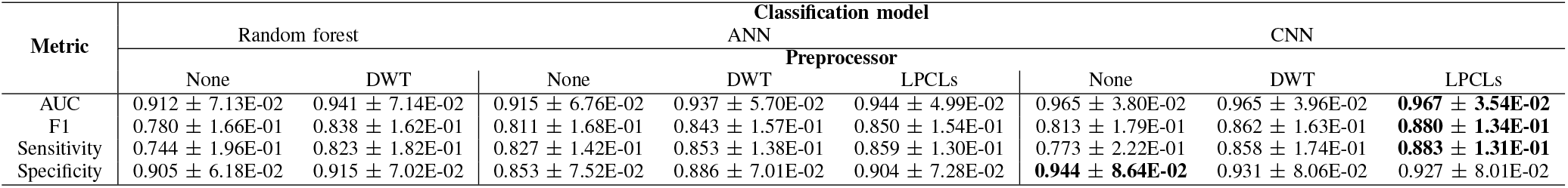
Numerical evaluation results. Total of 50 intersubject validations are conducted, with 30 independent runs per test patient data. Thus, a mean of 1,500 runs is calculated (Mean ± STD). The highest values for each metric are bolded. The individual values of the intersubject validations are given in Appendix B

**Fig. 5.**
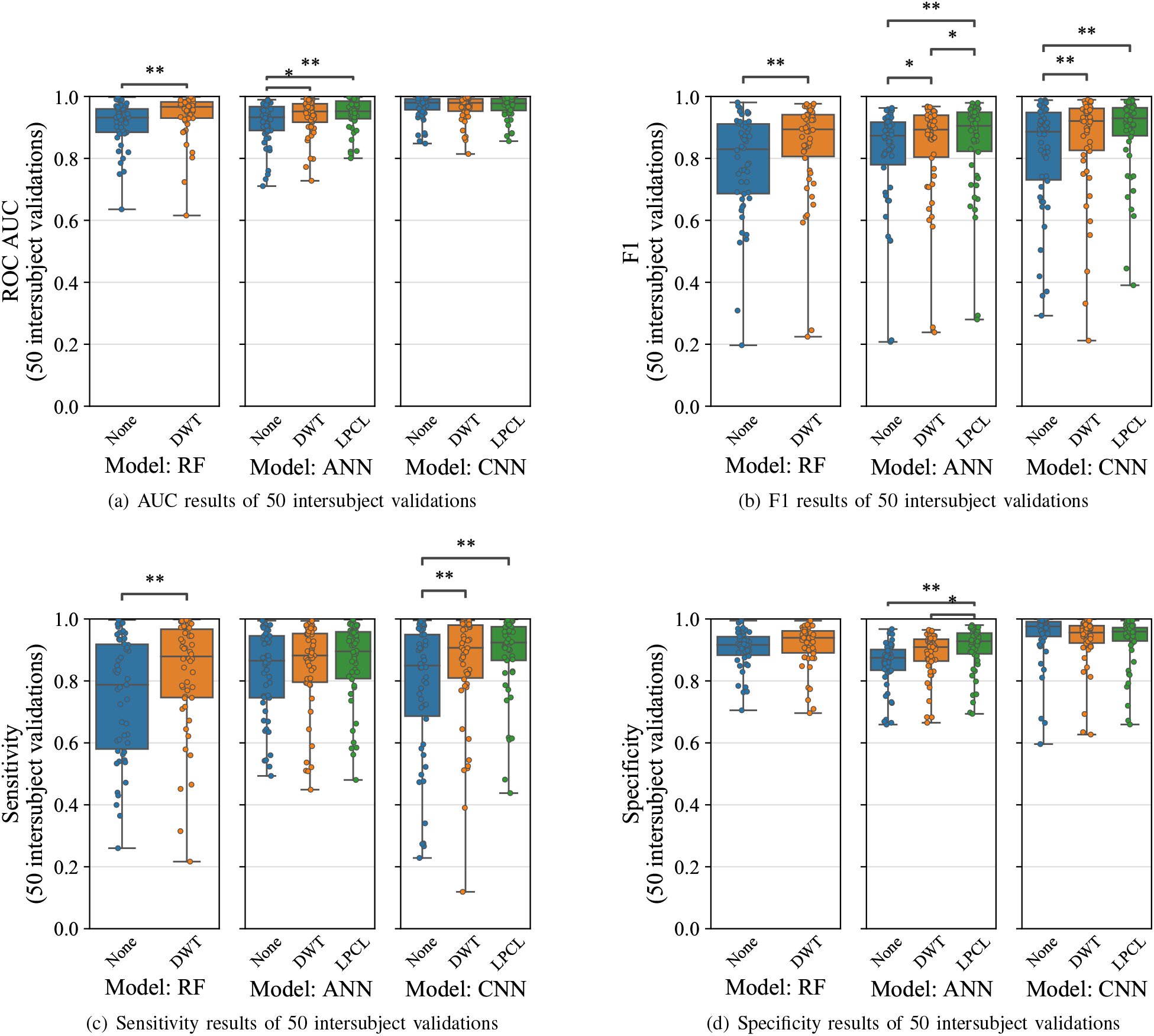
Visualized results in understanding the differences between preprocessors. Statistical significance is indicated by an asterisk (*: *p* < 0.05, **: *p* < 0.01).

Fig. 6 provides an example of prediction by CNN combined with the LPCL. In this figure, a relatively sharp waveform indicates an epileptic spike, regardless of its amplitude. Figs. 7 and 8 illustrate examples of the frequency responses at the proposed layers. In addition, Figs. 7 and 8 show clearly that the proposed method’s filter emphasizes the low-frequency band (around 12 Hz). Thus, while the conventional method manually focuses on the low-frequency band, it can be said that the proposed method automatically extracts this frequency. Moreover, Figs. 7(b) and 8(b) show that filters with odd symmetric constraints pass different frequency bands according to the filter length.

**Fig. 6.**
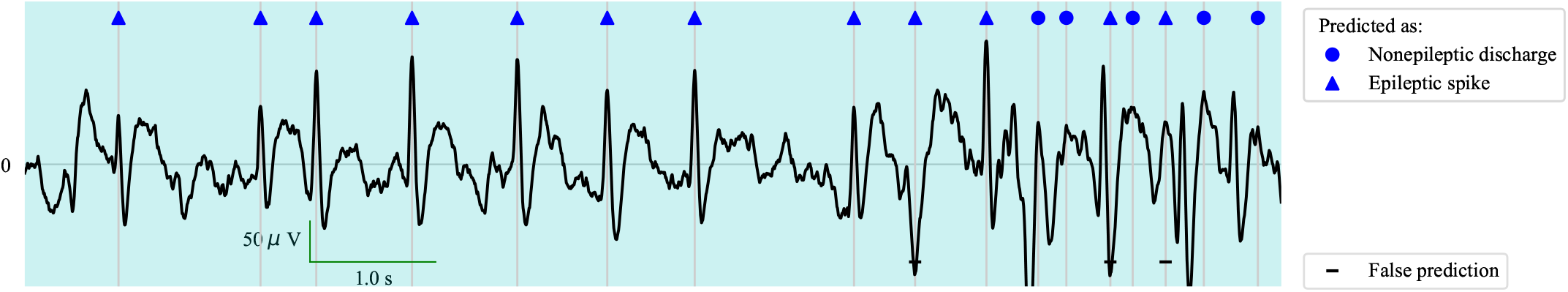
An example of the predicted spikes. The circles and triangles indicate nonepileptic discharges and epileptic spikes, respectively. The bars at the bottom indicate that the classification failed.

**Fig. 7.**
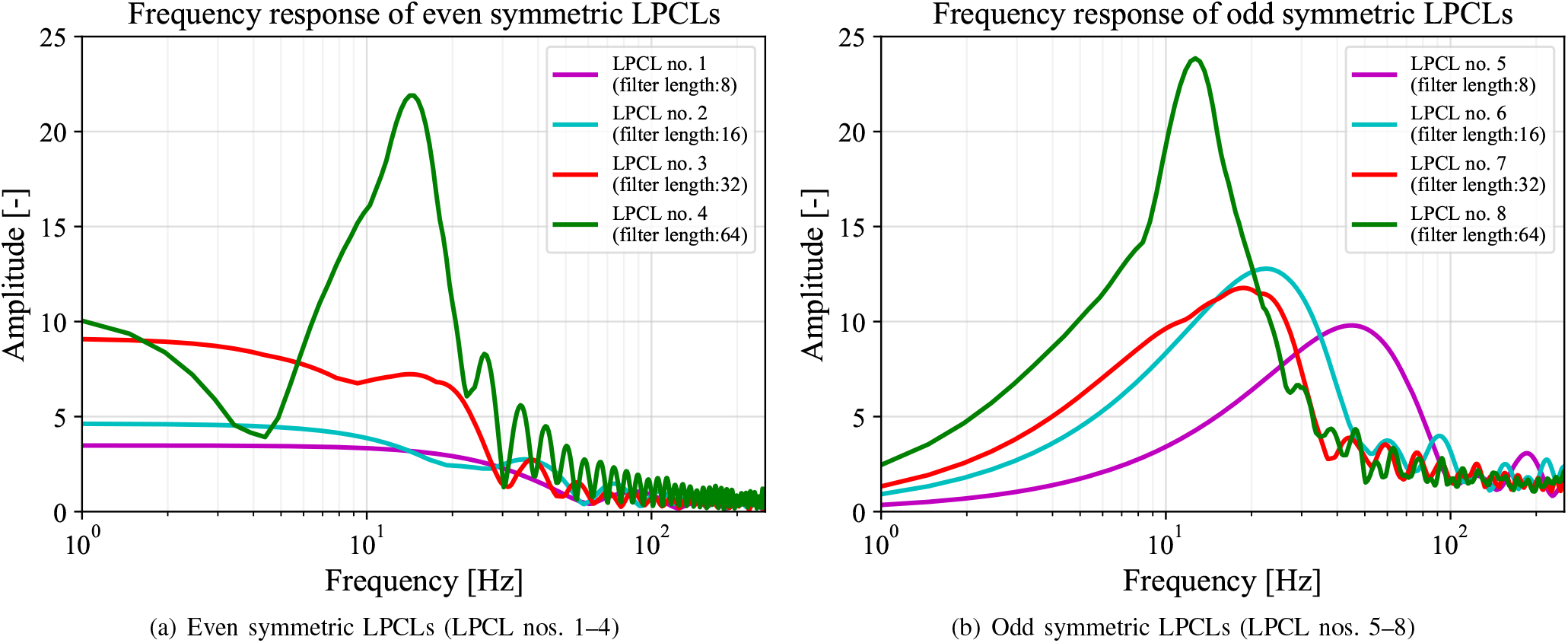
An example of mean filter spectrums a t the LPCL combining with ANN.

**Fig. 8.**
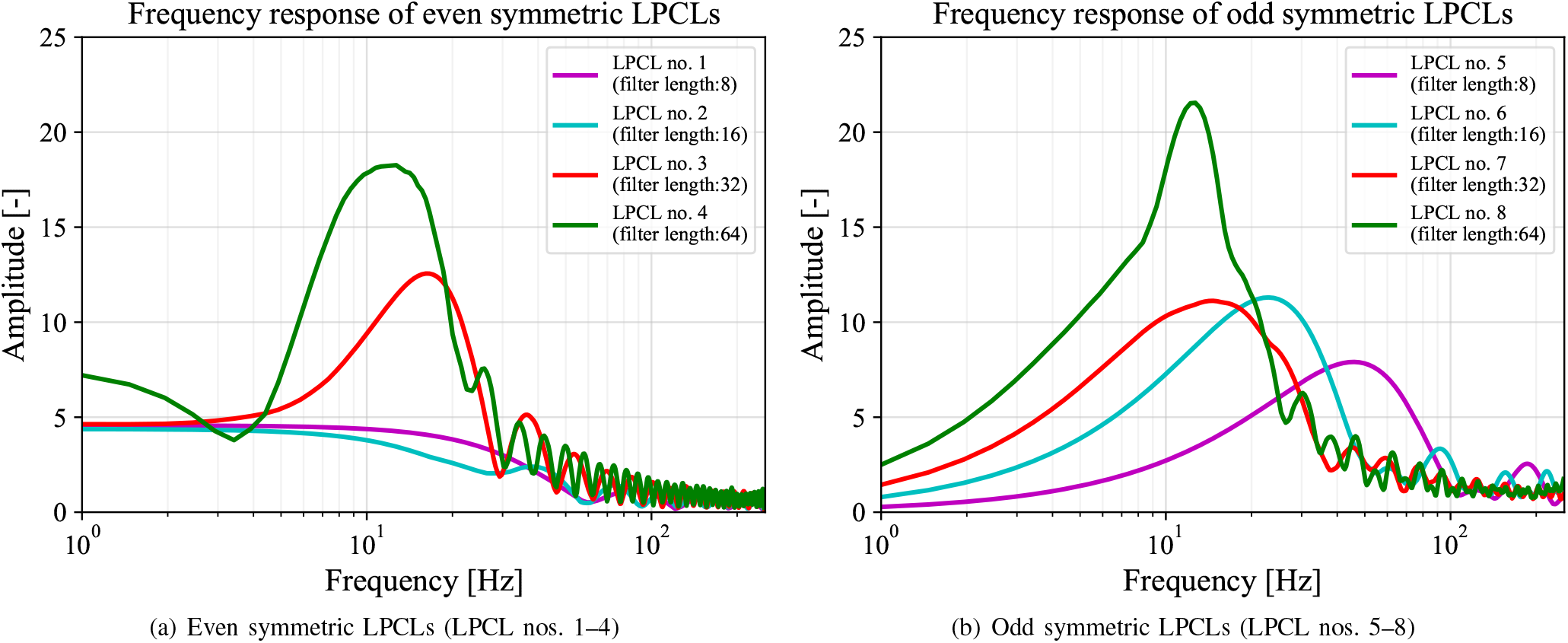
An example of mean filter spectrums a t the LPCL combining with CNN.

## V. Discussion and Conclusion

The experimental results show that the filters with an odd symmetry constraint have a different passband, as shown in Figs. 7 and 8. This behavior is similar to a bank of filters. As Fig. 8(b) shows, three of the frequency bands, approximately 12, 24, and 50 Hz (the focus bands of nos. 7 and 8 are similar), are focused on by the odd symmetry LPCLs. Focusing on the adjacent peak frequencies in the spectrum, the lower frequency is approximately half of the higher frequency. Their three frequency bands can be regarded as corresponding to the standard clinical frequency bands of *α*, *β*, and *γ*, respectively. This paper’s finding showed that the filters learned from the raw EEG and that the experts’ labels can be decomposed into the frequency bands contributing to the inspections. That is, the data-driven filters may emulate the logic of the physician’s analysis. Another advantage of the proposed work is that fine-tuning of the frequency bands is accomplished in a data-driven manner, such that the performance of the classifier is enhanced (AUC = 0.967, F1 = 0.880), as shown in Table V. Considering medical applications, the fact that the combination of LPCLs with CNN has achieved higher sensitivities than other methods [3], [8] is promising, as shown in Fig. 5(c).

Moreover, the LPCLs achieves these advantage points with a small computational complexity. As shown in (4) and (5), an LPCL consist of *L*/2-time additions (or subtractions) and an inner product calculation of size *L*/2. That is, only *L*/2 parameters (*h*_0_, *h*_1_, … , *h*_L/2−1_) are increased at the inner product calculation as learning parameters. In the proposed model shown in Fig. 4, there are a total of eight LPCLs with four different lengths (*L* = 8, 16, 32, and 64) and two constraint types, even and odd. In this case, the total number of parameters in the LPCLs is 120 only. Note that since the total number of parameters in the CNN-based model is approximately 3,500, the ratio of the number of parameters in the LPCLs to the all model’s parameters is less than 4%. Therefore, the ratio of the LPCL parameters in the overall architecture is relatively low. However, because the proposed method is designed based on neural networks, it cannot be combined with traditional classifiers like RF. In addition, similar to standard convolutional layers, it still requires a manual setting of hyperparameters such as the kernel size and number of filters. Considering these limitations, using DWT to decompose the EEG into clinical frequency bands [3], [28], [29] would prove to be better when it comes to versatility.

Next, we investigated the characteristics of the 1-s segments to consider the effectiveness of the frequency band extracted by the LPCLs. To determine the differences of spectra between the nonepileptic discharge segments and epileptic spike segments, statistical analyses were performed on the amplitude distributions at each frequency using Welch’s *t*-test [41]. Then, the effect sizes were calculated using Cohen’s *d* [42]. Fig. 9 shows the mean spectrum of all 15,004 nonepileptic discharges, the mean spectrum of all 15,833 epileptic spikes, the areas where *p* < 0.01 in the *t*-test, and the effect sizes. Fig. 9 shows that there are significant differences (*p* < 0.01) in the amplitudes of almost all frequencies. In addition, in the range of 5–15 Hz, there is a large difference (*d* ≈ 0.8) between the two classes. Similarly, the LPCLs, especially no. 8, as shown in Fig. 7(b), showed a strong response to this significantly different low-frequency band. This resulted in LPCLs that can extract the frequency bands of statistical interest in the proposed data-driven approach. Furthermore, because the methods using the LPCL and the predefined filter of 4–64 Hz exhibit comparable performance, as shown in Fig. 5, a frequency band such as those shown in Figs. 7 and 8—less than 30 Hz, as roughly estimated—rather than a much higher frequency band is sufficient for epileptic spike detection.

**Fig. 9.**
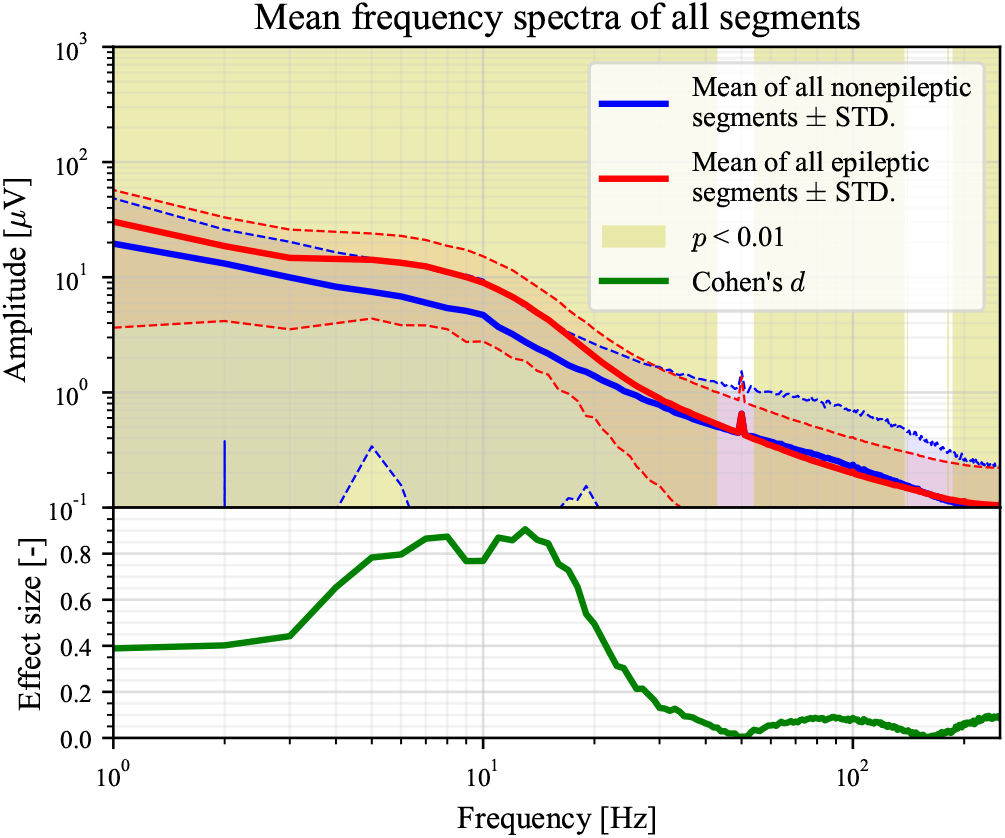
The mean spectrum of all 15,004 segments of nonepileptic discharges and the mean spectrum of all 15,833 segments of epileptic spikes. The areas where *p* < 0.01 in the *t*-test between the two classes at each frequency are filled in with yellow, and the bottom of the graph shows its effect size.

Finally, we consider the advantage of the dataset. In this paper, EEGs were measured from 50 CECTS patients, and 15,833 epileptic spikes and 15,004 nonepileptic discharges were then extracted as 1-s segments. To the best of our knowledge, the number of epileptic spike segments is the largest in the literature on epileptic spike detection, as described in Section II-C. This number of segments strongly supports the credibility of the statistical validation in this paper. However, more non-epileptic labels would be needed for the task of finding epileptic spikes in whole EEG recordings, rather than for the EEG segment classification task, as in this work. Moreover, the results of this paper may be limited by the fact that all patients’ symptoms are CECTS.

In the design of this dataset, we set the segment’s length as 1 s, following other studies [3], [23] and the annotation tasks performed by the five specialists. Of course, certain studies have used different length segments [14], [22]. As the results of this paper show, 1-s extraction is sufficient to achieve a high AUC (> 0.9 in most cases) for CECTS spikes. In particular, because epileptic spike-wave discharges in CECTS patients are known to contain a 3–4 Hz component [43], a segment length of 1 s can fully contain one of these discharges. Furthermore, even if the position of extracting the spike waveform is slightly misaligned, it is unlikely that any part of the waveform will be lost; thus, the 1-s extraction is appropriate.

In conclusion, we proposed a method to combine a bank of LP filters with a NN-based model and the ability to learn its coefficients from the data. To the best of our knowledge, we have built the largest dataset in the literature, containing 30,837 samples annotated by two neurosurgeons, one clinical technologists, and one pediatrician. The proposed model classifies 1-s segments as epileptic spikes or nonepileptic discharges with high performance (AUC > 0.9 in most cases). Furthermore, the filter’s frequency response fitted from the EEG is strong in the low-frequency range (around 12 Hz). This band coincided brilliantly with the frequency band of interest in the raw EEG segments of epileptic spikes.

## Acknowledgments

The authors would like to thank Ms. Meiko Sakurai and Ms. Junko Hirota for assistance with annotating, and Ms. Yuiko Kumagai for a fruitful discussion of statistical analysis.

## Appendix A EEG dataset

Table VI shows the dataset information used in the experiment. This dataset was labeled by two neurosurgeons, one clinical technologists, and one pediatrician. The total number of labeled samples is 30,837. The EEG recordings contain both awake and sleep—rapid eye movement (REM) or non-REM—states. This paper did not separate these states because each peak can be observed in both states.

**TABLE VI.** Dataset information of 50 Epileptic EEG records diagnosed with CECTS.

| Patient ID | Age of years | Sex | Recording duration |  | Annotator | #Epileptic spikes | #Nonepileptic discharges | #Total |
| --- | --- | --- | --- | --- | --- | --- | --- | --- |
| 1 | 7 | Female | 21 m | 52 s | Pediatrician | 424 | 178 | 602 |
| 2 | 6 | Male | 23 m | 53 s | Pediatrician | 263 | 629 | 892 |
| 3 | 5 | Male | 29 m | 45 s | Pediatrician | 240 | 580 | 820 |
| 4 | 10 | Female | 21 m | 50 s | Pediatrician | 236 | 461 | 697 |
| 5 | 6 | Female | 28 m | 38 s | Pediatrician | 939 | 728 | 1,667 |
| 6 | 7 | Male | 26 m | 48 s | Neurosurgeon 1 | 105 | 207 | 312 |
| 7 | 10 | Male | 28 m | 50 s | Clinical technologist | 321 | 351 | 672 |
| 8 | 6 | Female | 28 m | 24 s | Clinical technologist | 166 | 297 | 463 |
| 9 | 6 | Female | 31 m | 19 s | Neurosurgeon 2 | 412 | 94 | 506 |
| 10 | 8 | Male | 29 m | 8 s | Neurosurgeon 1 | 495 | 239 | 734 |
| 11 | 7 | Female | 28 m | 56 s | Neurosurgeon 1 | 345 | 199 | 544 |
| 12 | 8 | Female | 26 m | 56 s | Clinical technologist | 655 | 189 | 844 |
| 13 | 10 | Female | 27 m | 28 s | Neurosurgeon 1 | 341 | 303 | 644 |
| 14 | 10 | Male | 25 m | 41 s | Neurosurgeon 1 | 295 | 210 | 505 |
| 15 | 6 | Female | 29 m | 10 s | Neurosurgeon 1 | 159 | 355 | 514 |
| 16 | 11 | Female | 32 m | 42 s | Neurosurgeon 1 | 175 | 340 | 515 |
| 17 | 11 | Female | 27 m | 49 s | Neurosurgeon 1 | 295 | 335 | 630 |
| 18 | 10 | Male | 24 m | 24 s | Neurosurgeon 1 | 385 | 189 | 574 |
| 19 | 6 | Male | 28 m | 14 s | Neurosurgeon 1 | 232 | 289 | 521 |
| 20 | 7 | Male | 28 m | 57 s | Pediatrician | 74 | 118 | 192 |
| 21 | 11 | Female | 27 m | 17 s | Neurosurgeon 1 | 446 | 385 | 831 |
| 22 | 9 | Male | 24 m | 13 s | Neurosurgeon 1 | 231 | 330 | 561 |
| 23 | 7 | Female | 29 m | 15 s | Neurosurgeon 1 | 253 | 390 | 643 |
| 24 | 7 | Male | 29 m | 45 s | Neurosurgeon 1 | 350 | 190 | 540 |
| 25 | 10 | Male | 29 m | 29 s | Pediatrician | 17 | 238 | 255 |
| 26 | 9 | Male | 29 m | 22 s | Neurosurgeon 1 | 408 | 389 | 797 |
| 27 | 8 | Male | 29 m | 48 s | Neurosurgeon 1 | 294 | 292 | 586 |
| 28 | 8 | Male | 26 m | 47 s | Neurosurgeon 1 | 1,009 | 763 | 1,772 |
| 29 | 7 | Female | 28 m | 37 s | Neurosurgeon 1 | 416 | 293 | 709 |
| 30 | 9 | Female | 27 m | 9 s | Neurosurgeon 1 | 129 | 245 | 374 |
| 31 | 9 | Male | 27 m | 15 s | Clinical technologist | 282 | 112 | 394 |
| 32 | 7 | Female | 26 m | 53 s | Clinical technologist | 560 | 314 | 874 |
| 33 | 9 | Female | 13 m | 13 s | Clinical technologist | 331 | 214 | 545 |
| 34 | 9 | Male | 32 m | 27 s | Pediatrician | 21 | 253 | 274 |
| 35 | 12 | Female | 28 m | 11 s | Neurosurgeon 1 | 355 | 342 | 697 |
| 36 | 6 | Female | 28 m | 12 s | Clinical technologist | 440 | 244 | 684 |
| 37 | 9 | Female | 19 m | 40 s | Neurosurgeon 1 | 368 | 358 | 726 |
| 38 | 6 | Female | 30 m | 4 s | Pediatrician | 315 | 301 | 616 |
| 39 | 9 | Female | 26 m | 13 s | Pediatrician | 456 | 225 | 681 |
| 40 | 5 | Male | 28 m | 39 s | Pediatrician | 211 | 255 | 466 |
| 41 | 11 | Male | 32 m | 27 s | Pediatrician | 318 | 85 | 403 |
| 42 | 7 | Male | 27 m | 54 s | Pediatrician | 256 | 209 | 465 |
| 43 | 9 | Female | 30 m | 4 s | Pediatrician | 172 | 108 | 280 |
| 44 | 10 | Female | 26 m | 41 s | Pediatrician | 99 | 55 | 154 |
| 45 | 7 | Female | 29 m | 34 s | Pediatrician | 261 | 322 | 583 |
| 46 | 7 | Female | 30 m | 1 s | Clinical technologist | 72 | 369 | 441 |
| 47 | 10 | Male | 27 m | 8 s | Pediatrician | 287 | 509 | 796 |
| 48 | 3 | Male | 29 m | 8 s | Pediatrician | 271 | 511 | 782 |
| 49 | 6 | Male | 27 m | 50 s | Pediatrician | 351 | 151 | 502 |
| 50 | 5 | Male | 23 m | 14 s | Pediatrician | 297 | 261 | 558 |
| Total |  | 24 males |  |  | 21 by Neurosurgeon 1<br>1 by Neurosurgeon 2<br>8 by Clinical technologist | 15,833 | 15,004 | 30,837 |
|  |  | 26 females |  |  | 20 by Pediatrician |  |  |  |

## Appendix B Numerical results

Tables VII to X list the results of individual intersubject validations as AUC, F1, sensitivity, and specificity, respectively. Table V and Fig. 5 are created based on these tables.

**TABLE VII.**
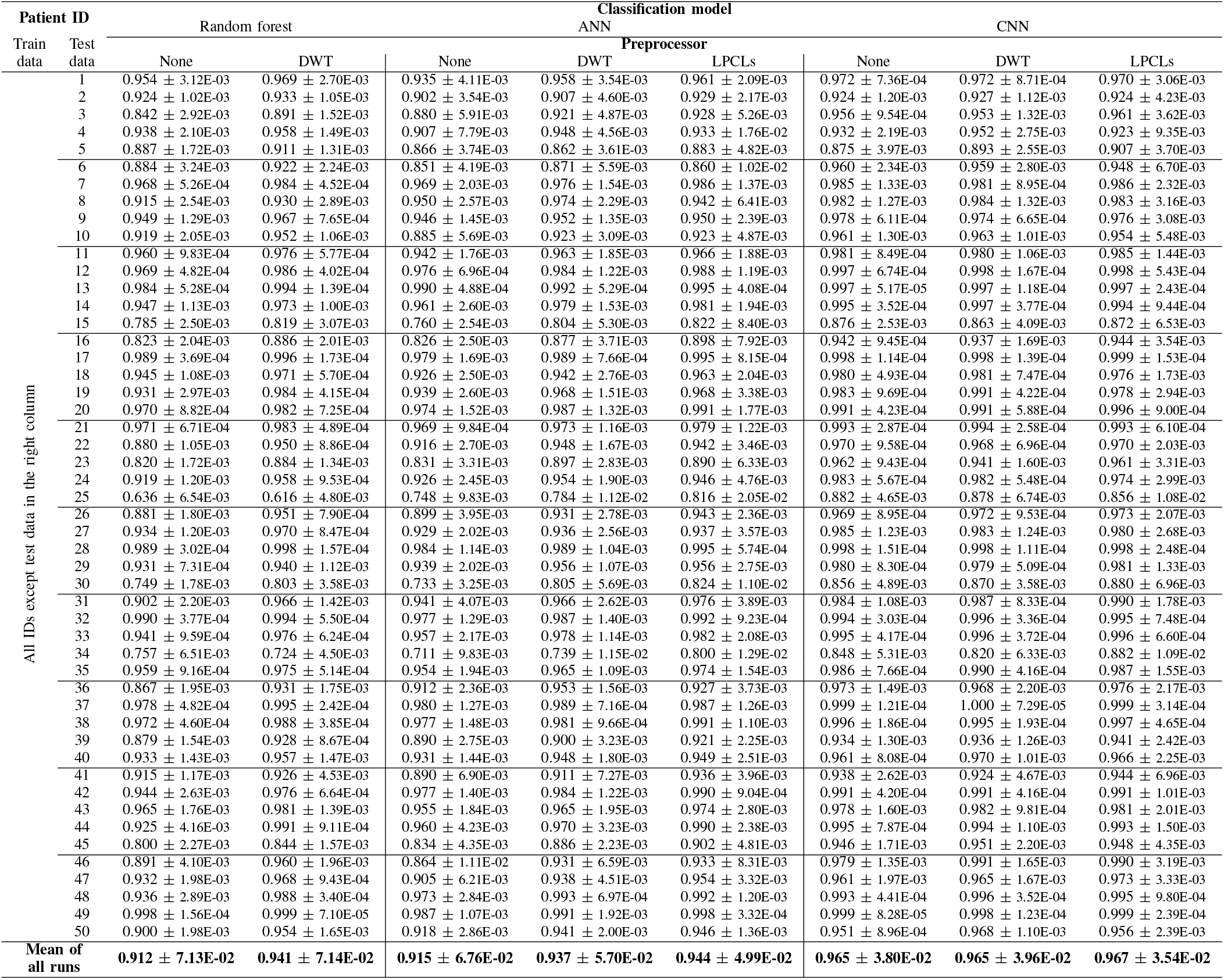
AUC evaluation results. For each method, the mean of 30 independent runs is calculated (Mean ± STD).

**TABLE VIII.**
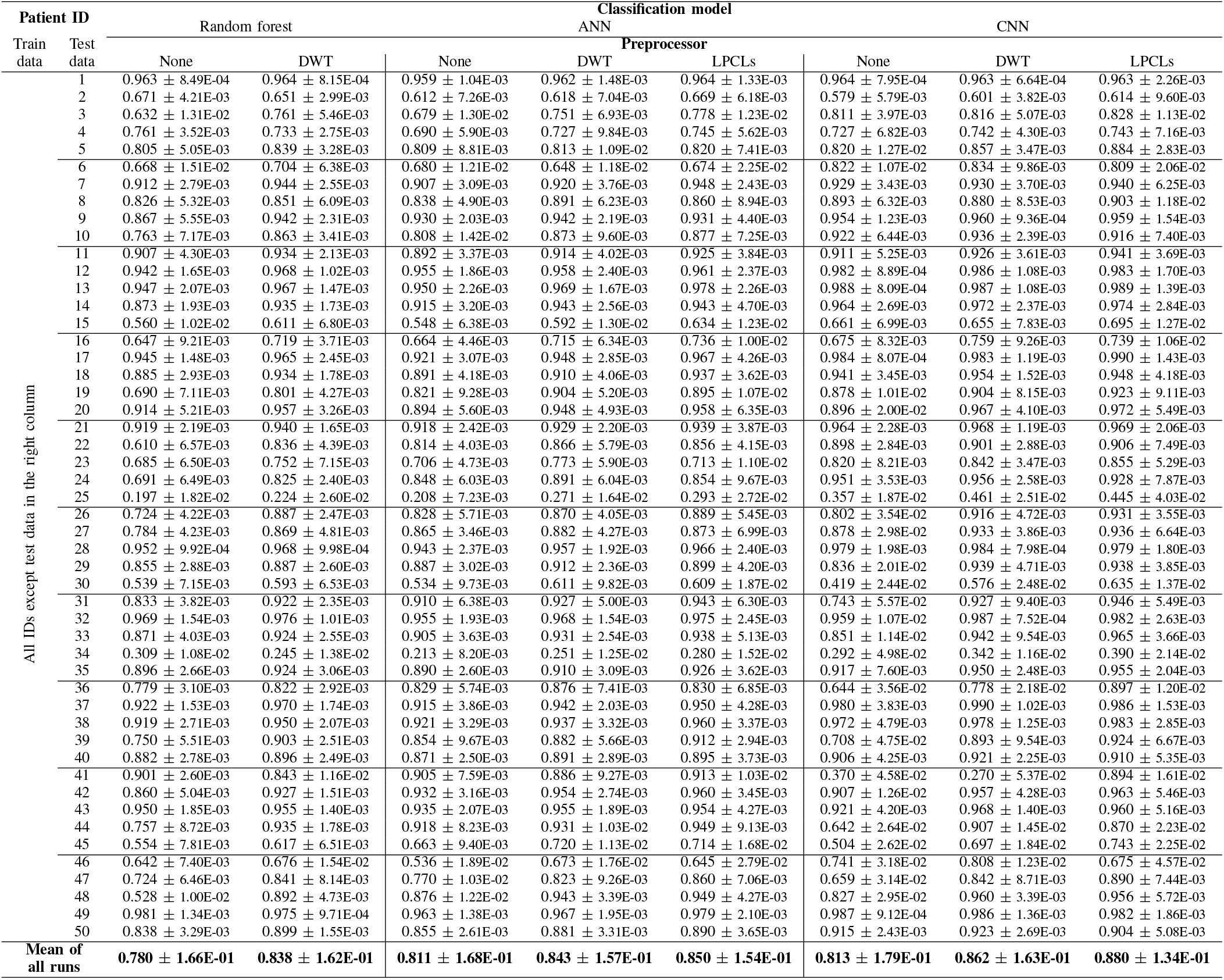
F1 value evaluation results. For each method, the mean of 30 independent runs is calculated (Mean ± STD).

**TABLE IX.**
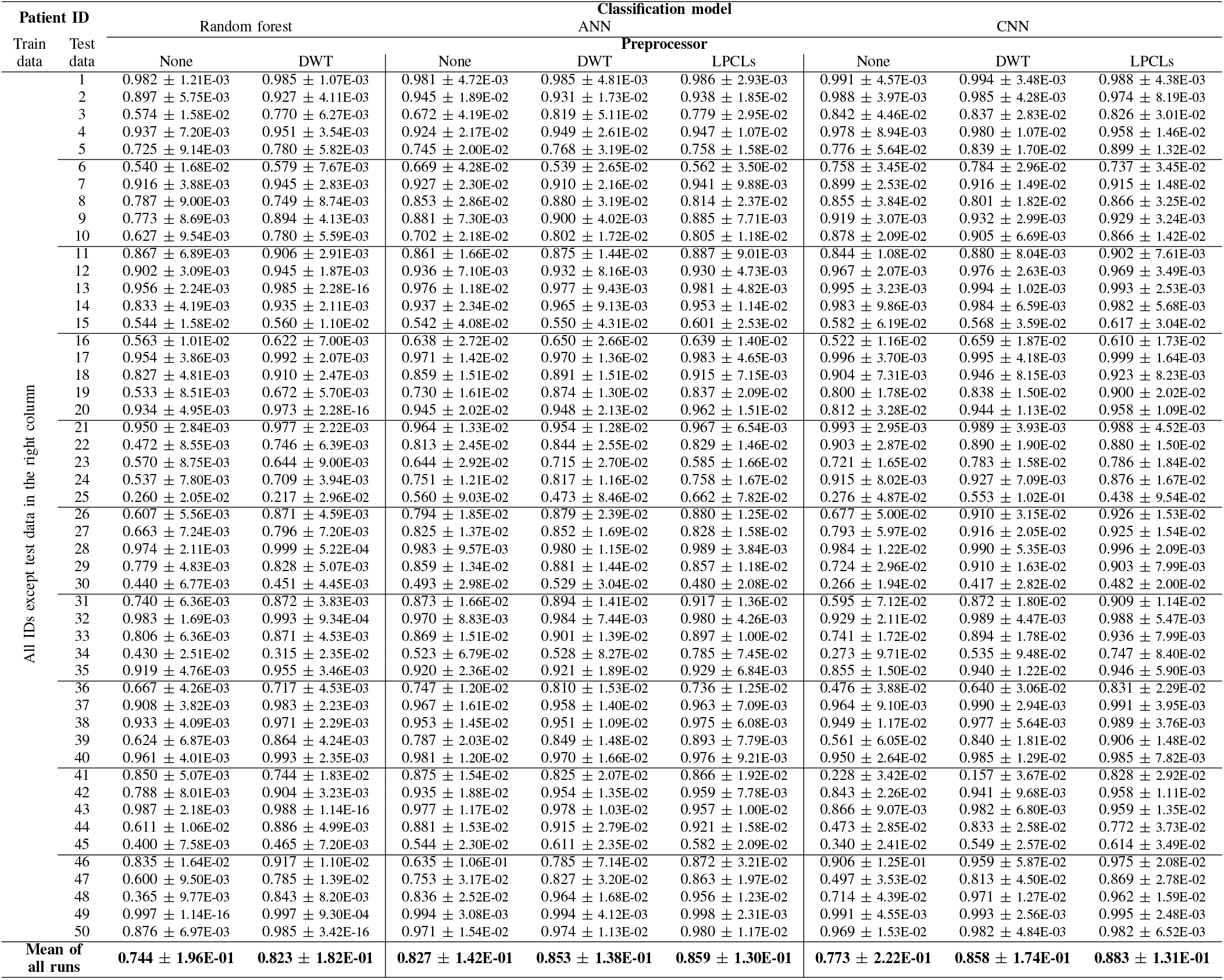
Sensitivity evaluation results. For each method, the mean of 30 independent runs is calculated (Mean ± STD).

**TABLE X.**
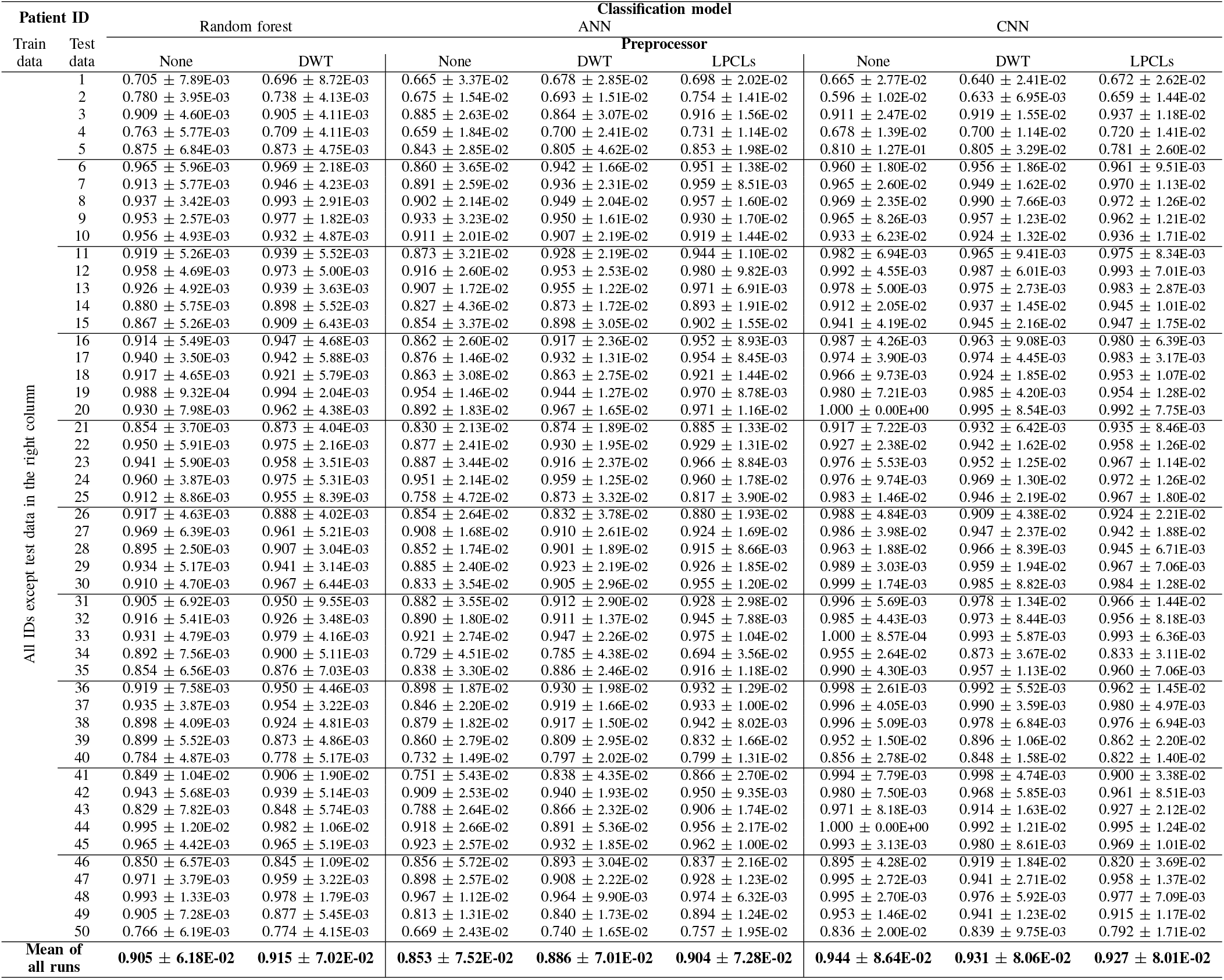
Specificity evaluation results. For each method, the mean of 30 independent runs is calculated (Mean ± STD).

